# Class I histone deacetylase complex: structure and functional correlates

**DOI:** 10.1101/2023.04.24.538028

**Authors:** Xiao Wang, Yannan Wang, Simiao Liu, Yi Zhang, Ke Xu, Liting Ji, Roger D. Kornberg, Heqiao Zhang

## Abstract

*Schizosaccharomyces pombe* Clr6S, a class I histone deacetylase complex, functions as a zinc-dependent enzyme to remove acetyl groups from lysine residues in histone tails. We report here the cryo-EM structures of Clr6S alone and in a complex with a nucleosome. The active center, revealed at near atomic resolution, includes features important for catalysis - a water molecule coordinated by zinc, the likely nucleophile for attack on the acetyl-lysine bond, and a loop that may position the substrate for catalysis. The cryo-EM map in the presence of a nucleosome reveals multiple Clr6S-nucleosome contacts, and a high degree of relative motion of Clr6S and the nucleosome. Such flexibility may be attributed to interaction at a site in the flexible histone tail, and is likely important for function of the deacetylase, which acts at multiple sites in other histone tails.

## INTRODUCTION

Post translational modifications (PTM) of the histone tails of nucleosomes influence transcriptional events^1–4^. Among PTMs, acetylation of lysine residues by histone acetyltransferase complexes (HATs)^5–7^ is associated with transcriptional activation, while deacetylation by histone deacetylases (HDACs) results in transcriptional repression^8, 9^. The opposing actions of HATs and HDACs maintain the balance of acetylation levels *in vivo*^10^.

There are four classes of HDACs in yeast and mammals. Class I (Rpd3-like) HDACs are zinc-dependent enzymes^8^, and the Rpd3 deacetylase of yeast occurs in two forms, large (Rpd3L) and small (Rpd3S) complexes^11–14^. Rpd3S, a 400 kDa complex of five subunits (Rpd3, Sin3, Ume1, Rco1, and Eaf3) functions downstream of the histone methyltransferase, SET2^15–17^. Histone H3 lysine 36 methylation (H3K36me) recruits Rpd3S to open reading frames, deacetylating the acetyl-lysines on histones H3 and H4, and thereby maintaining a hypoacetylation state of the regions. SET2-Rpd3S signaling is indispensable for suppressing cryptic transcription, which often occurs within open reading frames^15–17^. Rpd3S preferentially binds to a di-nucleosome, stimulating deacetylation of the two constituent mono-nucleosomes^18–20^. Previous studies have identified the counterparts of Rpd3S subunits in *Schizosaccharomyces pombe* (*S. pombe*): Pst2 (corresponding to Sin3), Clr6 (corresponding to Rpd3), Prw1 (corresponding to Ume1), Alp13 (corresponding to Eaf3), and Cph1 and Cph2 (corresponding to Rco1)^21–24^. These *S. pombe* proteins form an H3K36me-dependent histone deacetylase complex, Clr6S, confirmed to be the homolog of Rpd3S in *S. cerevisiae*^22, 23^.

Structures of HDAC-related complexes are limited to some single HDAC proteins^25–28^, a few subcomplexes^29–31^, and the class II HDAC complex^32^. The Rpd3S/Clr6S complex has been extensively studied for almost two decades, but structural information has not been available.

## RESULTS

### H3K36me3 binding affinity and deacetylase activities of Clr6S and Rpd3S

*S. cerevisiae* Rpd3S and its *S. pombe* homolog Clr6S were purified to virtual homogeneity (Fig. 1A). Both complexes exhibited micromolar binding affinity for a synthetic H3K36me3 peptide, measured by Surface Plasmon Resonance (Fig. 1B). We developed an HDAC assay with the use of a nucleosome bearing the H3K36me3 modification and acetylated at multiple sites in histone H4 (Fig. 1C). Western blot analyses with site-specific antibodies showed that Rpd3S and Clr6S possessed histone deacetylase activity toward H4K5Ac (acetylated Lys5), H4K8Ac (acetylated Lys8), H4K12Ac (acetylated Lys12), and H4K16Ac (acetylated Lys16) (Fig. 1D).

**Fig 1.**
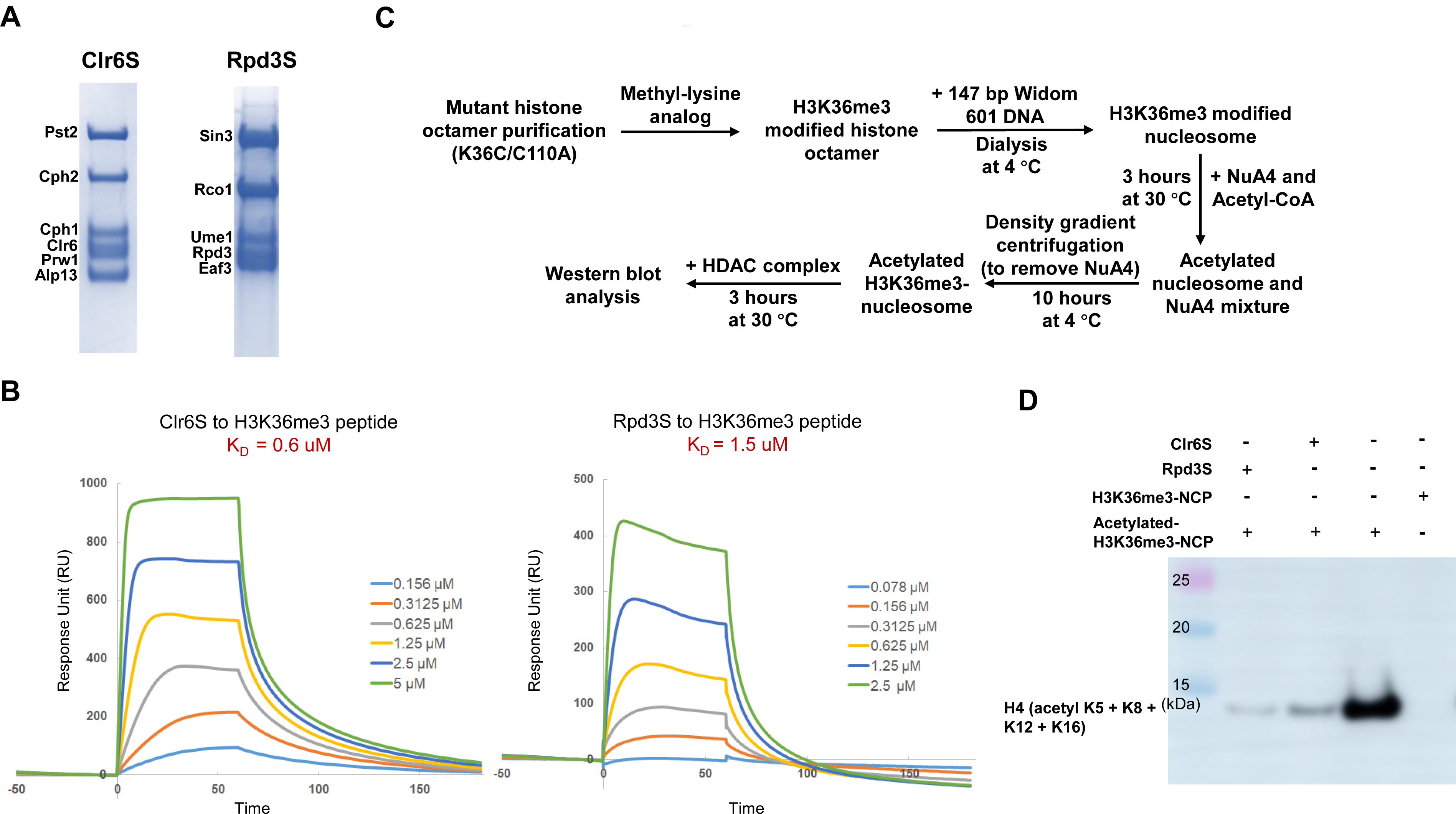
Biochemical analyses of the purified *S. pombe* Clr6S and *S. cerevisiae* Rpd3S complexes. (A) SDS-PAGE analyses of Clr6S and Rpd3S. The samples after size-exclusion chromatography were loaded on a 12% resolving gel and stained with Coomassie blue. (B) SPR analyses of Clr6S and Rpd3S binding to the synthesized H3K36me3 peptides. Binding curves at different concentrations are color-coded. (C) A workflow for the histone deacetylase assay of Clr6S, Rpd3S or Clr6S mutant protein on the acetylated-H4 substrate established in this study. (D) HDAC results of Clr6S and Rpd3S reacted with the acetylated-H3K36me3-NCP. Western blots were performed using anti-H4 (acetyl K5 +K8 +K12 +K16).

### Structure of the Clr6S complex

Rpd3S and Clr6S were subjected to cryo-EM analysis. Rpd3S particles embedded in vitreous ice exhibited preferred orientations, whereas Clr6S particles were suitable for structure determination (Fig. S1). The Clr6S complex was stabilized by GraFix (gradient fixation). Data collection and multiple rounds of processing yielded a 3.2 Å cryo-EM map (Fig. 2A and Fig. S2). A structure of Clr6S, built by a combination of homology-based and *de novo* modeling, was a good fit to the cryo-EM map (Fig. S3).

**Fig 2.**
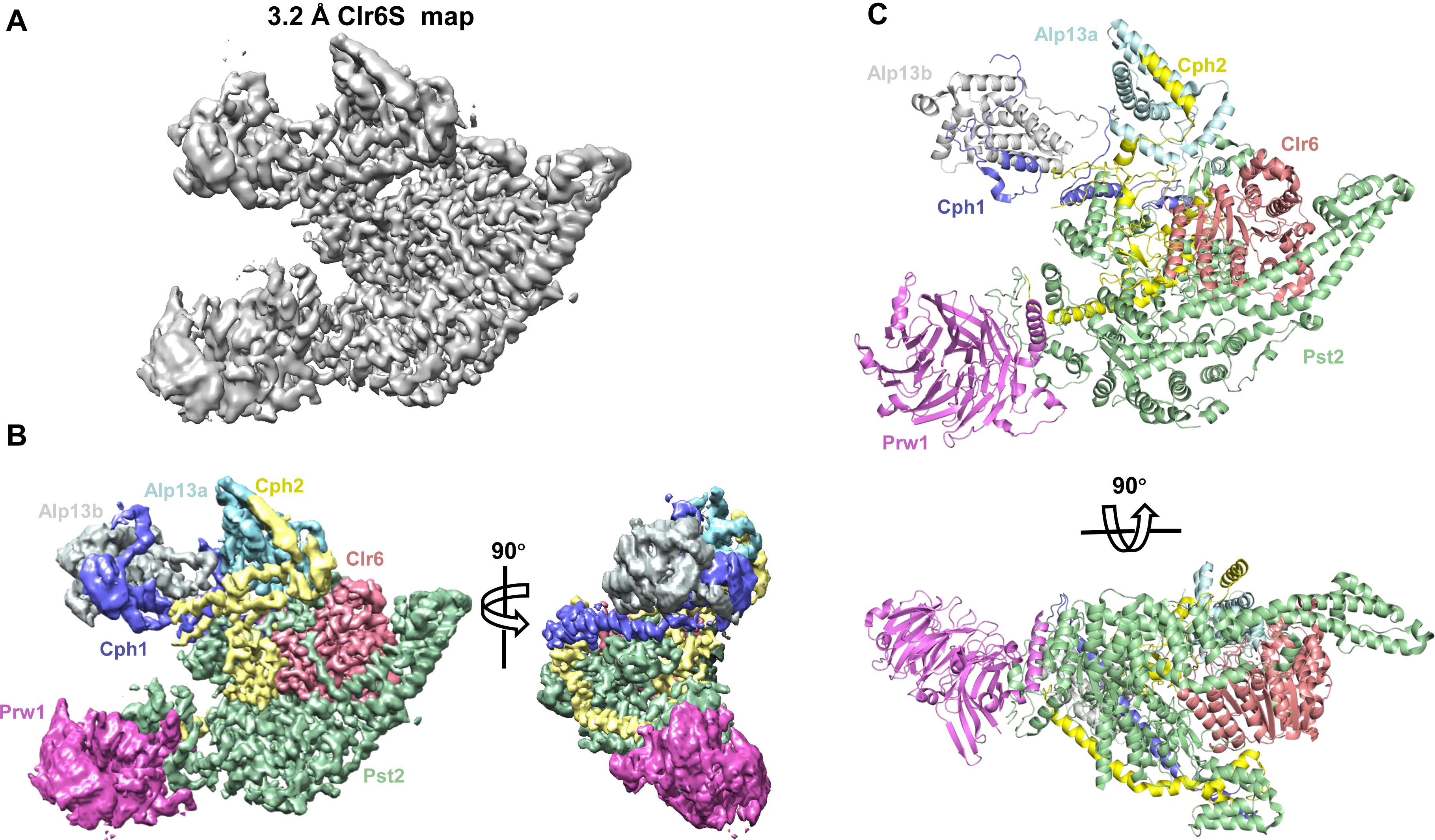
Overall structure of the Clr6S complex. (A) 3.2 Å cryo-EM map of Clr6S. (B) Segmentation of the Clr6S map. Different subunits are colored and labeled accordingly. (C) Ribbon diagrams of Clr6S structure, rotated by 90 °. Each subunit is colored and labeled aside.

All six subunits of the Clr6S complex are seen in the map (Fig. 2B, Fig. 2C and Table S1). Unexpectedly, the structure also reveals two copies of Alp13 (hereafter referred to as Alp13a and Alp13b) (Fig. 2C), an MRG (MORF4-Related Gene) family protein, whose stoichiometric information was absent from all previous genetic and biochemical studies, resulting in a seven subunit Clr6S complex. The structure shows that Pst2, Clr6, and Prw1, a WD40-containing subunit, form a subcomplex, within which Pst2 associates with the catalytic subunit Clr6, as well as interacting with Prw1 (Fig. 2C), consistent with a previous study showing that the *S. cerevisiae* counterparts Sin3, Rpd3 and Ume1 form a stable complex independent of other subunits^33^. Clr6 is embraced by Pst2, Alp13a, and Cph2, leaving the catalytic pocket accessible from only one side of the complex (Fig. 2C).

Alp13a associates with Cph2, while Alp13b is connected to the complex by both Cph1 and Cph2 (Fig. 2C and Fig. 3A). Alp13 consists of two domains, an N-terminal chromo domain, not visible in the cryo-EM map, and a C-terminal MRG domain. There is no apparent contact between the two Alp13 subunits. Although Alp13a and Alp13b contact the core complex in different ways (Fig. 3A and 3B), both copies of Alp13 interact with Cph1 and Cph2 through structurally conserved interfaces. The α3 helix of the MRG domain of Alp13b contacts an α-helix of Cph2, forming interface 1 (Fig. 3C). The next α-helix of Cph2 interacts with the α1 helix of Alp13b, forming interface 2. The similar pattern of interactions was observed for Alp13a-Cph1 (Fig. 3D), and also in the previously reported solution structure of Pf1-MRG15 (Fig. 3E)^34^, indicative of the conservation of this interaction pattern among the MRG domain-containing proteins and their binding partners.

**Fig 3.**
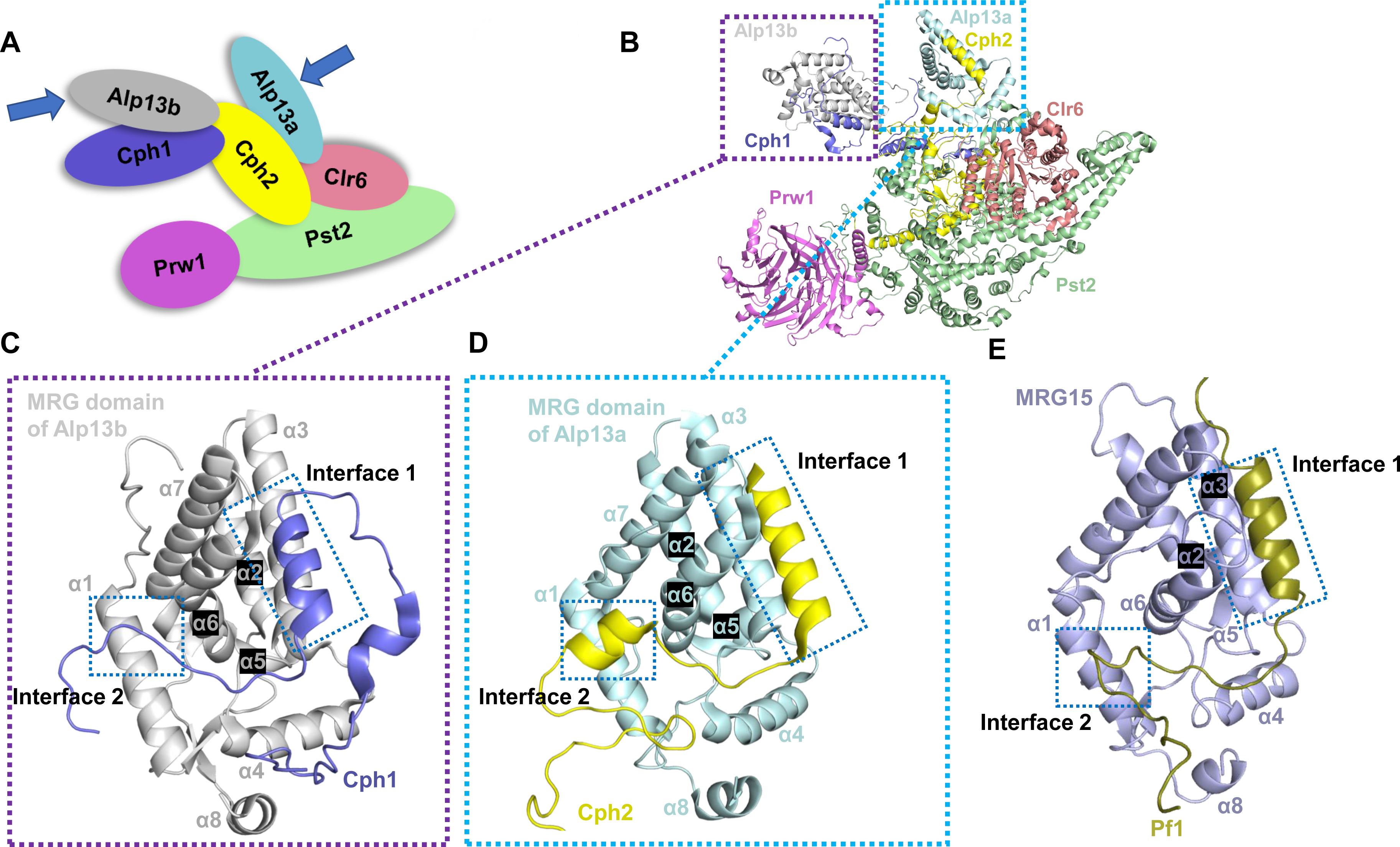
Two copies of Alp13 subunits in the Clr6S complex. (A) A scheme of the Clr6S complex. Each subunit is colored as Fig. 2C. Two copies of Alp13 are indicated with blue arrows. (B) Ribbon diagrams of Clr6S structure. The interfaces around Alp13a and Alp13b are highlighted with cyan and purple dashed boxes. (C)-(E) are the expanded views of the interaction details between the MRG domains of Alp13b, Alp13a and MRG15 and the Cph1, Cph2 and Pf-1, respectively. The interfaces are highlighted with blue dashed boxes.

### Scaffolding roles of Pst2 and Cph2

Pst2, like its *S. cerevisiae* homolog Sin3, is the largest subunit of the HDAC complex. The structure shows that Pst2 consists of five domains, termed PAH1 (paired amphipathic helix 1), PAH2 (paired amphipathic helix 2), PAH3 (paired amphipathic helix 3), ARM, and C-terminal domain (CTD) (Fig. 4A and 4B). PAH1 and PAH2 are in close proximity to the catalytic core of Clr6, though no direct interaction is apparent (Fig. 4C). PAH3 is sandwiched between a plant homeodomain (PHD) of Cph2 and a long C-terminal α helix of Cph1. A solution structure (PDB entry: 2LD7) of a binary complex of SAP30, a subunit of the mammalian Rpd3L/Sin3A histone deacetylase complex, with PAH3 of Sin3A, revealed a similar interaction^32^, involving several aromatic amino acids and leucines from PAH3 and a phenylalanine in the interacting helix of Cph1 or SAP30, which are highly conserved (Fig. 4C). The ARM region of Pst2 forms extensive contacts with Clr6, mainly through hydrogen bond interactions. The CTD domain of Pst2 interacts with the N-terminal α-helix and the last two blades of the WD40 protein, Prw1 (Fig. 4C). Pst2 evidently serves as a structural platform, bringing together almost all the other subunits of the complex.

**Fig 4.**
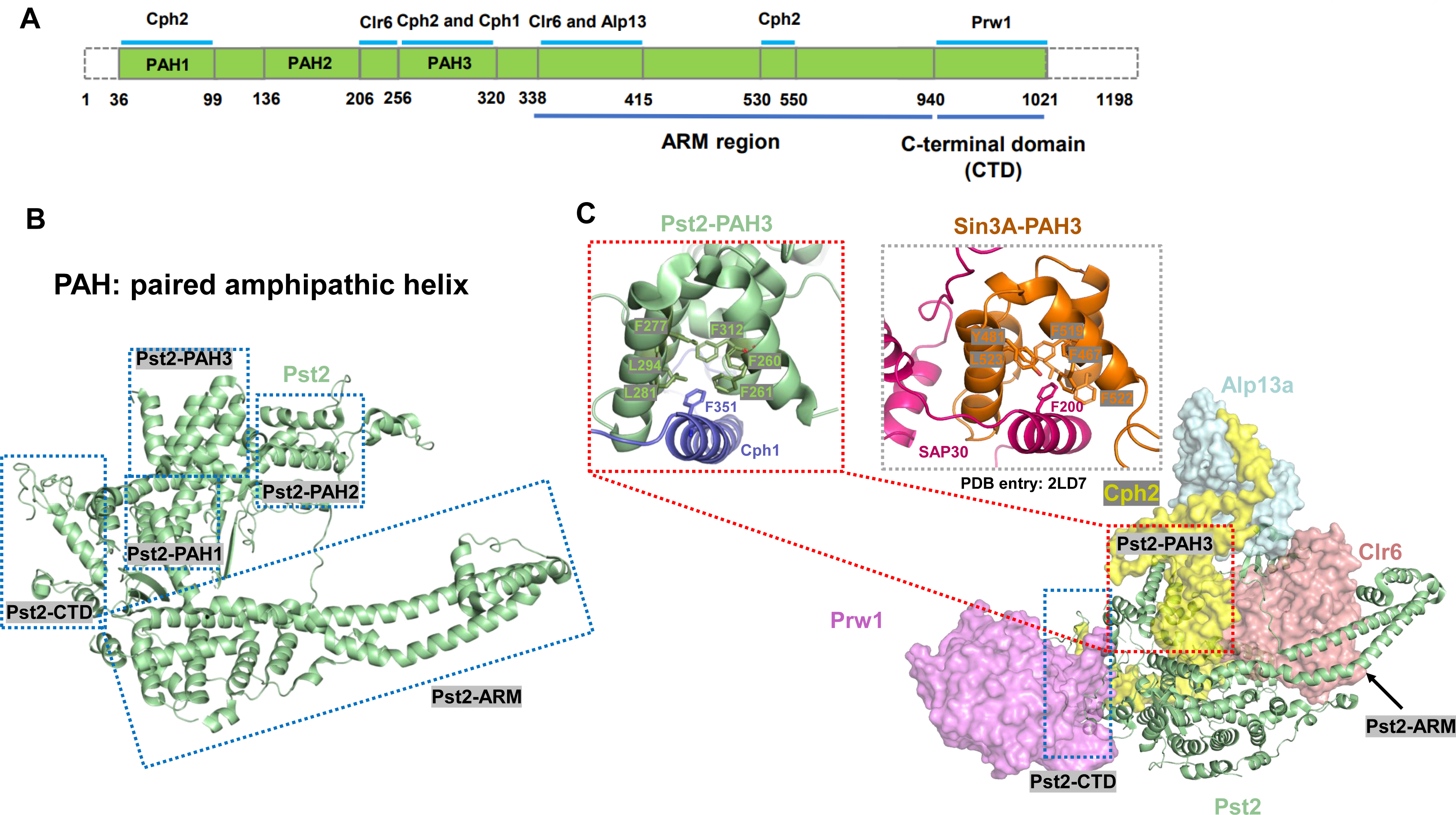
Pst2 serves as a structural platform for protein complex assembly. (A) Domain organization of the Pst2 subunit. The boundary residues of each domain are labeled on the bottom. The interacting regions of Pst2 with other subunits are labeled on the top. The PAH1, PAH2, PAH3, ARM and CTD domains are labeled. (B). Ribbon diagram of the Pst2 subunit. Five domains of Pst2 are indicated with blue dashed boxes. (C) The interaction details of Pst2 with other subunits. The interaction between Pst2-PAH3 and Cph2 is shown in a close-up view. The residues contributed to the interaction are shown in stick. Single-letter abbreviations for the amino acids shown in the figure are as follows: F, Phe; L, Leu; Y, Tyr.

Cph2 contains two PHD domains, the first of which is not visible in the structure. The visible region (residues 333 to 599) adopts an extended conformation (Fig. 2C), centered on the second PHD domain, with a region before, the Inter-PHD domain, and a region after, the post-PHD domain (Fig. 5A). The cryo-EM map contained density consistent with two zinc ions in the second PHD domain, coordinated by Cys409, Cys412, His433 and Cys436, and by Cys425, Cys428, Cys453 and His456. The Inter-PHD domain interacts with the MRG domain of Alp13a. The second PHD domain lies between the PAH1 and PAH3 domains of Pst2 and Clr6 (Fig. 5B). The post-PHD domain bridges the Cph1-Alp13b heterodimer and the PAH3 domain of Pst2. Thus Cph2 interacts with all the other subunits except Prw1, whose association with the complex occurs only through the Pst2-CTD (Fig. 4C).

**Fig 5.**
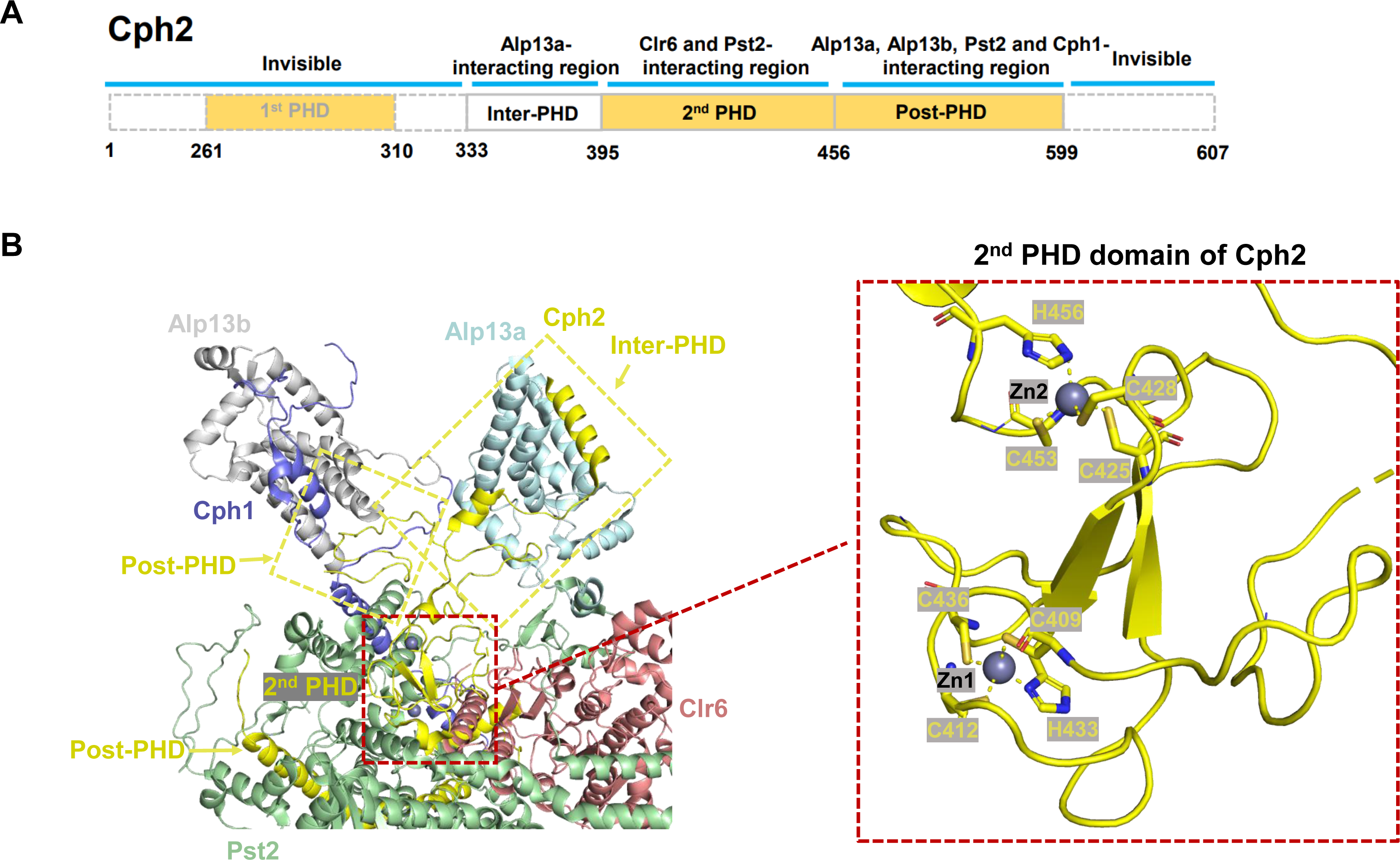
Scaffolding role of Cph2. (A) Domain organization of Cph2. The boundary residues of each domain are indicated on the bottom. (B) From left to right are the Cph2 structure and its interactions with other subunits in the complex, and an expanded view of its PHD domain. The zinc ions are shown as gray spheres. The residues constituting the two zinc-binding motifs are shown in stick and labeled. Single-letter abbreviations for the amino acids shown in the figure are as follows: C, Cys; H, His.

### Functional correlates: conserved active center loop, active center coordination chemistry

The Clr6 region of the cryo-EM map was at higher resolution than other parts, revealing the catalytic pocket in near atomic detail. The map contained density for a metal ion, presumably a zinc ion, and a water molecule, anticipated from previous studies^26, 27^ (Fig. 6A, 6B, and 6C). The putative zinc ion is coordinated by Asp173, His175, and Asp261, and by the water molecule. The water molecule is additionally coordinated by His137 and His138 (Fig. 6B and 6C). Sequence alignment shows that these and other residues, and a tyrosine, Tyr300, are highly conserved in the HDAC family, likely due to their significance in catalysis (Fig. 6D). Previous structural study of the single human HDAC protein indicates that the Tyr300 residue located in the pocket may be important for substrate binding and catalysis^26, 27, 30^. The zinc ion and the conserved residues in multiple HDAC structures superimpose well upon one another (Fig. 6E).

**Fig 6.**
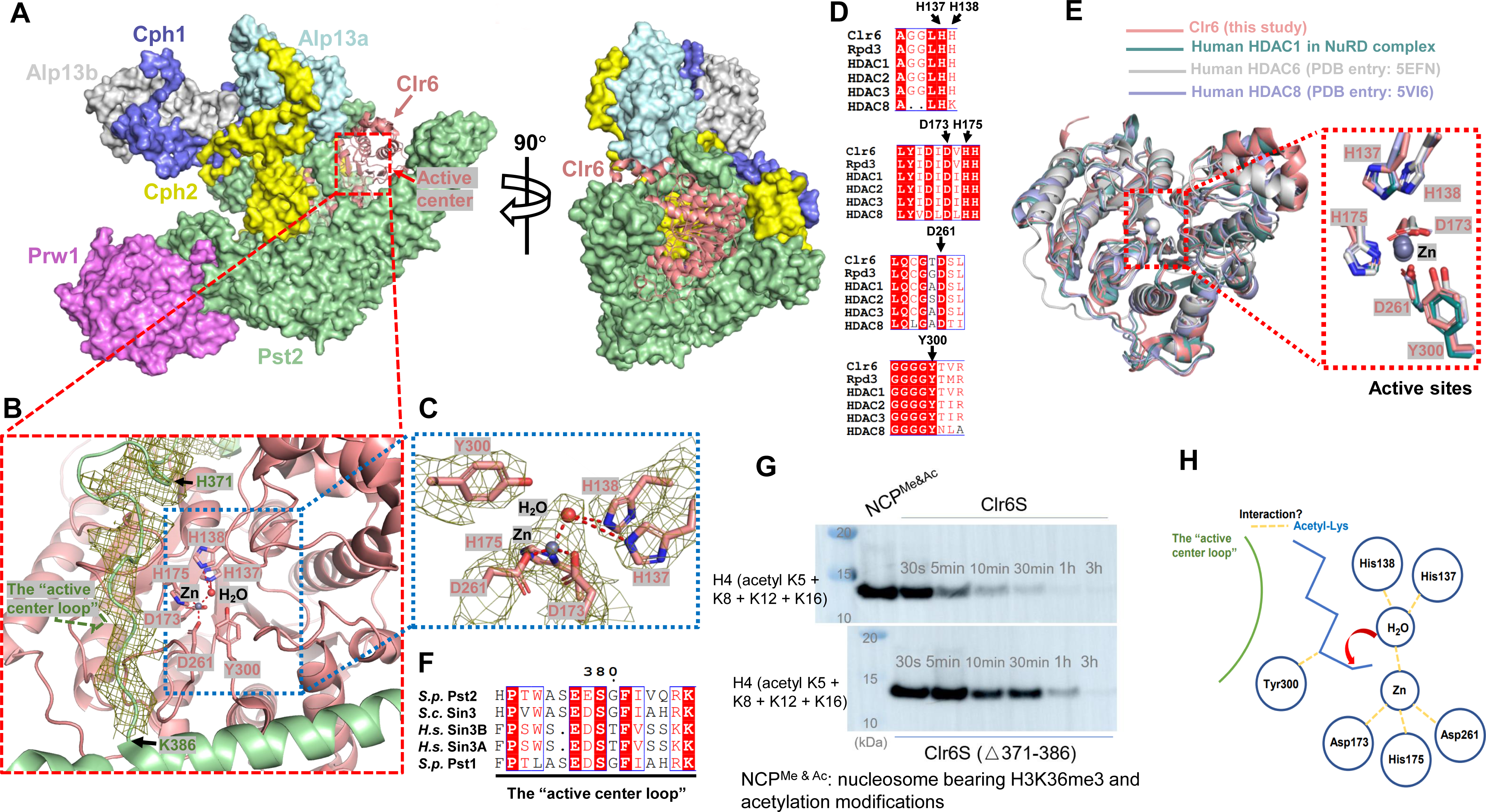
Active center of the Clr6S complex. (A) The catalytic subunit Clr6 in the complex. Clr6 is shown as cartoon, and other subunits are shown as surface. The active center in Clr6 is highlighted with a red dashed box. (B) A close-up view of the active center of Clr6S complex. The active sites are shown in stick, and the “active center loop” of Pst2 is shown as mesh and labeled. Gray and red spheres represent the zinc ion and water molecule located in the active center of Clr6, respectively. Single-letter abbreviations for the amino acids shown in the figure are as follows: D, Asp; H, His; Y, Tyr. (C) A close-up view of the zinc ion and the water molecule bound in the active center. The electron density for the putative zinc ion, the water molecule and their coordinating residues are contoured at 4.5σ. (D) Structure superposition of the Clr6 and its active sites with other HDAC proteins. The active sites are shown in stick. (E) Sequence alignments of the residues located in the active center. The residues important for catalysis are indicated with black arrows. (F) Sequence alignment of the “active center loop” of Pst2 with other Sin3-family members. *S.p.* represents *Schizosaccharomyces pombe*, *H.s.* represents *Homo sapiens*, and *S.c.* represents *Saccharomyces cerevisiae*. (G) Comparison of HDAC activity of wild-type Clr6S and Clr6S (△371-386) proteins. (H) A proposed model illustrating the catalytic mechanism. The catalytic residues are shown as blue balls, and the polar interactions are indicated with yellow dashed lines.

A loop (residues 371-386) of Pst2 adjacent to Clr6 near the active center (Fig. 6B) is highly conserved among Sin3-family members, including *S. cerevisiae* Sin3, *Homo sapiens* Sin3A and Sin3B, and *S. pombe* Pst1 (Fig. 6F). To investigate the possible importance of this “active center loop” for the deacetylase activity of Clr6S, it was deleted from Pst2, and a truncated form of Clr6S (Clr6S (△371-386)) was purified to homogeneity for measurement of HDAC activity (Fig. S4). Deletion of the active center loop diminished HDAC activity by an order of magnitude (Fig. 6G), consistent with a role of the loop in catalytic activity, although a secondary effect, due to perturbation of the protein structure could not be excluded.

These findings support a catalytic mechanism in which the zinc ion, Tyr300, and the “active center loop” accommodate the substrate, positioning it for catalysis, and the water molecule serves as a nucleophile for attack on the acetyl-lysine bond (Fig. 6H).

### Clr6S-nucleosome complex

The interaction of Clr6S with an H3K36me-modified, acetylated nucleosome (Fig. 1C) was investigated by electrophoretic mobility shift analysis (EMSA), and evidence for a Clr6S-nucleosome complex was obtained (Fig. 7A). The complex was stabilized by GraFix (Fig. S5A) and subjected to cryo-EM analysis (Fig. 7B, Fig. S5B and S5C). A 3.9 Å electron density map was obtained (Fig. 7C 7D, 7E and Fig. S6), which displayed extra density on the concave side of Clr6S, the location of the second PHD domain of Cph2 (Fig. 7C and 7D). The size and shape of the extra density were consistent with a nucleosome, but structural details of the nucleosome were not observed, indicative of motion relative to Clr6S. The density was roughly spherical, suggestive of rotational motion (Fig. 7E). We also obtained a cryo-EM map of an Rpd3S-nucleosome complex (Fig. 7F and Fig. S7), in which the HDAC complex was less well resolved but the nucleosome structure was clearly apparent (Fig. 7F). The location of the nucleosome was the same as that of the extra density in the Clr6S-nucleosome map, confirming the attribution of the extra density to a nucleosome.

**Fig 7.**
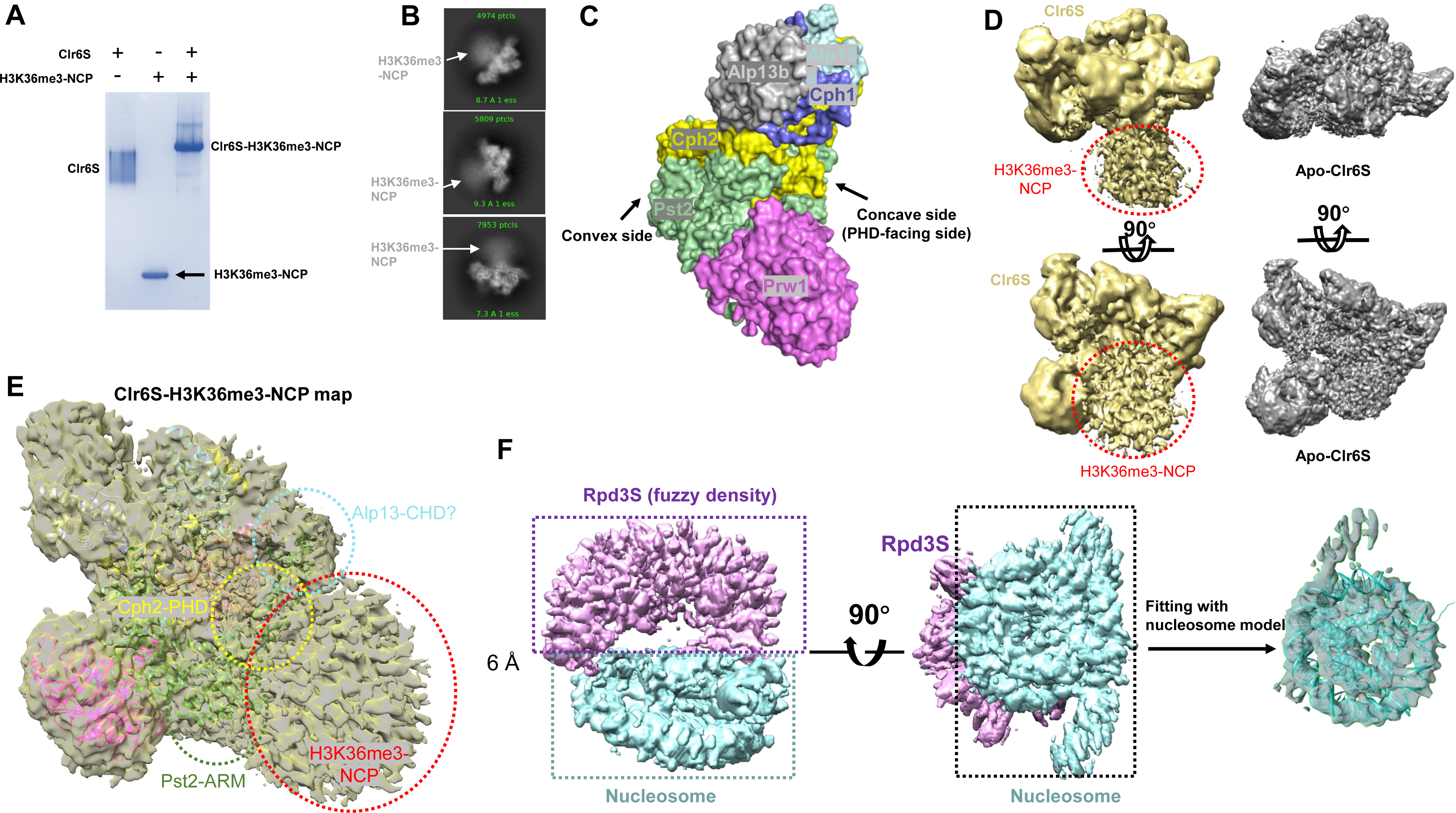
Interaction of Clr6S complex and the nucleosome. (A) The gel-shift result of Clr6S with H3K36me3-NCP, visualized by Coomassie blue staining. (B) 2D class averages of the Clr6S-H3K36me3-NCP. The nucleosome is indicated with a gray arrow. (C) Surface view of the Clr6S complex. The concave and convex sides of Clr6S are indicated with black arrows. (D) Comparison of the cryo-EM maps for Apo-Clr6S and Clr6S-H3K36me3-NCP. The maps of Apo-Clr6S and Clr6S-H3K36me3-NCP are shown in gray and yellow, respectively. (E) The binding surfaces of H3K36me3-NCP on Clr6S. Clr6S model was docked into the Clr6S-H3K36me3-NCP map as a rigid body. Clr6S subunits are shown as cartoon and colored as in Fig. 2C. The map transparency is adjusted to 50%. The interfaces of nucleosome-Alp13-CHD, nucleosome-Cph2-PHD, and nucleosome-Pst2-ARM are highlighted with cyan, yellow, and green dashed circles, respectively. (F) A 6 Å cryo-EM map of Rpd3S-H3K36me3-NCP, and rotated by 90 degree. The nucleosome is highlighted with a cyan dashed box, and the Rpd3S is highlighted with a purple dashed box. The figure was generated using Chimera. On the right is a fit of a nucleosome model (generated from PDB-7EA5) to the EM density. The nucleosome model is shown as cartoon representation.

## DISCUSSION

MRG domain-containing proteins, such as Alp13, occur in many chromatin regulatory complexes. However, multiple copies of an MRG domain-containing protein in a single complex, such as we find for the Clr6S complex, has not been previously reported. The preference of Rpd3S/Clr6S for di-nucleosomes raises the question whether the two chromodomains of Alp13a and Alp13b are responsible for interaction with the two constituent mono-nucleosomes.

Despite the low resolution of the nucleosome in the Clr6S-nucleosome complex, interactions with several Clr6S subunits are apparent. Additional poorly resolved density protruding from the MRG domain of Alp13a, absent from the density map of Clr6S alone, may be attributed to the chromo domain of Alp13a (Alp13-CHD, highlighted with a cyan dashed circle in Fig. 7E). Contacts of the PHD domain of Cph2 (Cph2-PHD) and the ARM domain of Pst2 (Pst2-ARM) with the nucleosome are also observed (Fig. 7E). These contacts are in keeping with previous work demonstrating the requirement for combined action of the chromo domain of Eaf3 and the PHD domain of Rco1 required for Rpd3S binding to nucleosomes^17^. The contacts may confer specificity for the H3K36me3-modified nucleosome and contribute to positioning the nucleosome for catalysis.

Both Clr6S-nucleosome and Rpd3S-nucleosome maps reveal flexibility of the HDAC-nucleosome interaction, but in a manner inversely related to one another. In the case of the Clr6S-nucleosome complex, the refinement is dominated by Clr6S and density due to the nucleosome is less well resolved, whereas in the case of the Rpd3S-nucleosome complex, the refinement is dominated by the nucleosome and density due to Rpd3S is less well resolved. Flexibility of the HDAC-nucleosome interaction is expected because the HDAC complex binds to H3K36, within the flexible histone tail. Flexibility may be necessary for the HDAC complex to act at multiple sites of acetylation elsewhere in the histone tails.

Previous studies of the HDAC1-MTA1 heterodimer, a component of the human NuRD complex, revealed that a D-myo-inositol-1,4,5,6-tetrakisphosphate (Ins(1,4,5,6)P_4_) molecule mediates the interaction of HDAC1 and MTA1 and enhances HDAC activity^30^. Comparison of our Clr6S structure with that of the previously reported HDAC1-MTA1 complex revealed steric clashes between MTA1 and Pst2 (Fig. S8A). Most importantly, no extra electron density was observed from the corresponding region of the Clr6S complex (Fig. S8B), suggesting that Clr6S may not require Ins(1,4,5,6)P_4_ as a cofactor. In contrast to the positively charged surface of the Ins(1,4,5,6)P_4_ binding pocket in HDAC1-MTA1 (Fig. S8C), the corresponding region in the Clr6S complex is apparently formed by a few negatively charged residues, in combination with other types of amino acids (Fig. S8D). Taken together, our structural analyses reveal the structural difference between the binding partners of class I HDACs and exclude the possibility of Ins(1,4,5,6)P_4_ binding to the Clr6S complex.

## Author contributions

H.Z. designed the experiments; X.W., and Y.W. carried out the experiments.; X.W., and H.Z. collected and processed the cryo-EM data.; H.Z. built the models.; H.Z. and R.D.K. wrote the paper with input from all authors.

## Data availability

Cryo-EM maps for the Apo-Clr6S complex and Clr6S-H3K36me3-NCP have been deposited in the Electron Microscopy Data Bank (EMDB) under accession codes EMD-35416 and EMD-35417, respectively. Coordinate file of Apo-Clr6S has been deposited in the Protein Data Bank (PDB) under accession number 8IFG.

## Acknowledgements

We thank the Bio-Electron Microscopy Facility of ShanghaiTech University for assistance in data collection. The research was supported by a grant from ShanghaiTech University, and a grant from the Shanxi Key Laboratory of Protein Structure Determination (202104010910006) to H.Z.

## Materials and Methods

### Purification of the Clr6 and Rpd3S complexes

The insect-cell expression system was used to purify Clr6S, Rpd3S and their mutant proteins. The genes encoding all the subunits of Clr6S, including Pst2, Clr6, Prw1, Alp13, Cph1, and Cph2, synthesized from GenScript Biotech, were cloned into pFastBac1 vector, respectively. Pst2 bears a Flag tag at its N-terminus to facilitate subsequent affinity purification. Each baculovirus was produced using the Bac-to-Bac baculovirus expression system (Invitrogen). The Clr6S complex was co-expressed in High-five cells by adding equal volume of the six baculovirus for 48 hours at 27 °C. After infection, one liter of High Five cells (2.0 × 10^6^ cells ml^−1^ cultured in ESF921 medium) was harvested, centrifuged for 10 minutes at 4 °C (3,000 rpm). The Cell pellets were resuspended with buffer containing 20 mM Tris, pH 8.0, 150 mM NaCl and 3 mM DTT, and lysed by sonication. The lysate was further cleared by high-speed centrifugation, purified using anti-Flag resins (GenScript Biotech), and further purified using Heparin and size-exclusion chromatography (Superose 6, GE Healthcare) in buffer containing 20 mM HEPES, pH 8.0, 100 mM NaCl and 3 mM DTT. The pooled peak fractions were concentrated to ∼3 mg/ml before use.

To obtain the Clr6S (△371-386) protein, Pst2 (residues 1-370) and Pst2 (residues 387-1075) were constructed respectively. The baculovirus for each construct was produced using the standard Bac-to-Bac baculovirus expression system (Invitrogen). Pst2 (residues 1-370) and Pst2 (residues 387-1075) were co-expressed with other five subunits in High Five cells. The protein was purified following the same procedures used to purify the wild-type Clr6S, as described above.

To obtain the *S. cerevisiae* Rpd3S complex, the genes encoding Sin3, bearing a Flag tag at its N terminus, Rpd3, Ume1, Rco1, and Eaf3, all synthesized from GenScript Biotech, were constructed into pFastBac1 vector, respectively. The overexpression and purification of Rpd3S were performed following the same procedures used to purify the wild-type Clr6S, as described above.

The *Xenopus* histone octamer was overexpressed, purified as previously described^35^. The 147 bp Widom-601 DNA was used to reconstitute nucleosomes as described. The DNA sequence is shown as follows:

ATCGAGAATCCCGGTGCCGAGGCCGCTCAATTGGTCGTAGACAGCTCTAGCACCGCTTAA ACGCACGTACGCGCTGTCCCCCGCGTTTTAACCGCCAAGGGGATTACTCCCTAGTCTCCAGGCACGTGTCAGATATATACATCCTGAT

### Cryo-EM grid preparation, data collection and image processing

**Apo-Clr6S:** a total of 150 μg Clr6S protein was loaded onto a linear glycerol gradient (10% to 30%) in dialysis buffer (20 mM HEPES, pH 8.0, 100 mM NaCl, 3 mM DTT) supplemented with 0-0.125% EM-grade glutaraldehyde for fixation, and then subjected to ultracentrifugation (40,000 rpm, 4 °C, 10 hours). Gradients were fractionated manually in 500 μl. The crosslinking was quenched by the addition of Tris (pH 8.0) solution to a final concentration of 40 mM. All fractions were analyzed by SDS-PAGE, and the peak fractions were pooled, dialyzed against the dialysis buffer containing 20 mM HEPES, pH 8.0, 100 mM NaCl and 3 mM DTT. The crosslinked sample was then concentrated to approximately 0.6 mg/ml before grid preparation.

**Clr6S-H3K36me3-NCP:** the purified Clr6S and an equal molar of H3K36me3-NCP were incubated for 1 hour on ice. The mixture was applied to a 10%–30% glycerol gradient in dialysis buffer (20 mM HEPES, pH 8.0, 100 mM NaCl, 3 mM DTT) supplemented with 0-0.125% glutaraldehyde for fixation, and then ultracentrifuged at 40,000 rpm for 10 hours at 4 °C in a Beckman SW41 Ti rotor. Gradients were manually fractionated in 500 μl, quenched by the addition of a final concentration of 40 mM Tris (pH 8.0), and then analyzed by SDS-PAGE. The peak fractions containing the Clr6S-H3K36me3-NCP were pooled and dialyzed against the dialysis buffer. The dialyzed sample was concentrated to approximately 1 mg/ml before grid preparation.

**Rpd3S-H3K36me3-NCP:** the purified Rpd3S complex was inculcated with an equal molar of H3K36me3-NCP for 1 hour on ice. The complex was then crosslinked by GraFix (as described above). The sample was concentrated to approximately 1 mg/ml before grid preparation.

Vitrobot Mark IV was used for grid plunging. 200-mesh Quantifoil R1.2/1.3 holey carbon grids were glow-discharged for 40 seconds before grid freezing. 3 μl of crosslinked Clr6S, Clr6S-H3K36me3-nucleosome, and Rpd3S-H3K36me3-nucleosome were applied to the grids, respectively. After incubation for 20 seconds, each grid was blotted using filter paper (Ted Pella Inc.) for 2.5 seconds and plunged into liquid ethane with 100% chamber humidity at 8 °C. All grids were stored in liquid nitrogen until data collection.

### Cryo-EM data collection and image processing

Automated data acquisitions were performed using SerialEM^36^ on Titan Krios equipped with a Gatan K3 Summit direct electron detector (Gatan, Inc.) and operating at 300 kV with a nominal magnification of 22,500 × and a pixel size of 0.53 Å (super-resolution mode). The data collection and processing details for apo-Clr6S are described as follows: a total of 4,158 images were automatically recorded in super-resolution mode, with a defocus range from 1.3 to 2.5 μm. Each movie stack was dose-fractionated to 40 frames with a total electron dose of ∼60 e^-^/Å^2^ and a total exposure time of ∼3.4 seconds. Movie stacks were motion-corrected using Patch Motion Correction, and the defocus value was estimated using Patch CTF estimation in cryoSPARC^37^. The particles were first automatically picked using Blob picker and filtered using Inspect Picks in cryoSPARC^37^. The picked particles were extracted with a box size of 600 pixels (bin=2), followed by 4 rounds of 2D classification, generating a template for subsequent template-based picking. A total of 3,161,808 particles picked from Template picker were extracted with a box size of 600 pixels (bin=2), and then subjected to 2 rounds of 2D classification. A total of 1,680,440 particles were selected and subjected to *ab initio* reconstructions (number of classes=6), followed by 2 rounds of heterogeneous refinement. Non-uniform (NU) refinement was performed for the best class selected from the last round of heterogeneous refinement, yielding a reconstruction of Clr6S at 3.2 Å.

**Clr6S-H3K36me3-NCP**: the dataset for Clr6S-H3K36me3-NCP was recorded on the same Titan Krios using the same parameters as Apo-Clr6S. A total of 3,864 images were automatically recorded in, with a defocus range from 1.3 to 2.0 μm. Motion correction and CTF estimation were performed using Patch Motion Correction and Patch CTF estimation in cryoSPARC^37^, respectively. The automatically picked particles using Blob picker were extracted with a box size of 600 pixels (bin=2), followed by 5 rounds of 2D classification, giving rise to a template for subsequent template-based picking. A total of 1,698,670 particles picked from Template picker were re-extracted with a box size of 600 pixels (bin=2), and then subjected to 3 rounds of 2D classification. A total of 195,291 particles were subjected to *ab initio* reconstruction (number of classes=3), followed by heterogeneous refinement. Non-uniform (NU) refinement was performed for the best class, yielding a reconstruction of Clr6S-H3K36me3-NCP at 3.9 Å. Local refinement, focused classification, and multibody refinement didn’t work in this case.

**Rpd3S-H3K36me3-NCP**: the dataset for Rpd3S-H3K36me3-NCP was recorded on the same Titan Krios using the same parameters as Apo-Clr6S. A total of 3,345 images were automatically recorded in, with a defocus range from 1.3 to 2.0 μm. Motion correction, CTF estimation, and auto picking of particles were performed in cryoSPARC^37^ as described for the Clr6S-H3K36me3-NCP sample. 2D classification, *ab initio* reconstruction, and heterogeneous refinement were performed in cryoSPARC^37^. A final round of NU-refinement resulted in a 6 Å map. Local refinement, focused classification, and multibody refinement didn’t work in this case.

### Model building

All the model buildings were performed using the Cryo EM module of Phenix package^38^, assisted by Chimera^39^, and COOT^40^. The AlphaFold predicted models of Pst2, Clr6, Prw1, Alp13, Cph1 and Cph2 were edited in PyMOL to keep accordance with the cryo-EM map. After editing, all the models were docked into the cryo-EM map and manually adjusted in COOT^40^, and then subjected to Phenix^38^ for several rounds of real-space refinement. All the structure figures were generated by PyMOL (https://pymol.org/2/) and Chimera^39^.

### SPR assay

The binding affinities of wild-type Clr6S and Rpd3S proteins to the synthesized H3K36me3 peptide (ATKAARKSAPATGGVK_36_(me3)KPHRYRPG) (GenScript Biotech) were determined using BIAcore T200 system (GE Healthcare) performed at 25 °C. The Clr6S and Rpd3S proteins were immobilized onto the series S CM5 sensor chips (Cytiva), respectively. Gradient concentrations of H3K36me3 for Clr6S from 156 nM to 5 μM and Rpd3S from 78 nM to 2.5 μM were flowed over the series S CM5 chip surface in running buffer (20 mM HEPES, pH 8.0, 150 mM NaCl, 3 mM DTT, and 0.005% [v/v] P20). The background signals from the immobilized protein with running buffer at corresponding concentrations were recorded as control, subsequently subtracted from the raw data for correction. The affinity values were generated, and the binding affinities were obtained using a 1:1 (Langmuir) binding fit model with Biacore T200 Evaluation software version 3.2 (GE Healthcare).

### Electrophoretic mobility shift assay

To test the interaction, Clr6S and H3K36me3-modified nucleosome were mixed at an equal molar and incubated for 30 minutes on ice, and then were loaded onto a 3%-12% Native-PAGE gel (Thermo Fisher Scientific) and stained with Coomassie blue.

### Histone H3 lysine 36 tri-methylation modification

The histone octamer bearing K36C and C110A mutations on H3 was purified as previously described^35^. Tri-methylation of the H3K36 was performed using Methyl-lysine Analog (MLA) as described^41^. The H3K36me3 modification was confirmed by Western bot analysis using anti-H3K36me3 antibody (ab9050, Abcam). The modified nucleosomes were stored at 4 °C.

### Histone deacetylase assay (HDAC assay)

To obtain acetylated nucleosome for HDAC assays, H3K36me3 modified nucleosome was inculcated with the NuA4 acetyltransferase complex at a molar ratio of 50:1. The histone acetyltransferase (HAT) assay was performed as previously described^35^, and the NuA4 complex was removed by density gradient centrifugation after reaction. Peak fractions containing the acetylated nucleosomes were pooled and concentrated for HDAC assay. All the HDAC reactions mentioned in this study were performed in a reaction buffer containing 20 mM HEPES (pH 8.0), 100 mM NaCl, and 5 mM MgCl_2_. 0.1 μM different HDAC complex (wild-type Clr6S, Clr6S (△371-386) or wild-type Rpd3S) and 1 μM acetylated nucleosomes were added to the reaction buffer, and incubated at 30 °C for 3 hours, respectively. For evaluation of the HDAC activity, each reaction mixture was taken at different time points including 30 seconds, 5 minutes, 10 minutes, 30 minutes, 1 hour, and 3 hours, and quenched by adding 4X sodium dodecyl sulfate (SDS) loading buffer and boiled for 10 minutes. The reaction mixtures were loaded to a 4%-20% SDS-PAGE gel (Thermo Fisher Scientific), and subsequently analyzed by Western blot using anti-histone H4 (acetyl K5 + K8 + K12 + K16) antibody (ab177790, Abcam). The signals were quantified by ChemiDoc (Bio-Rad).

---

**Fig S1.**
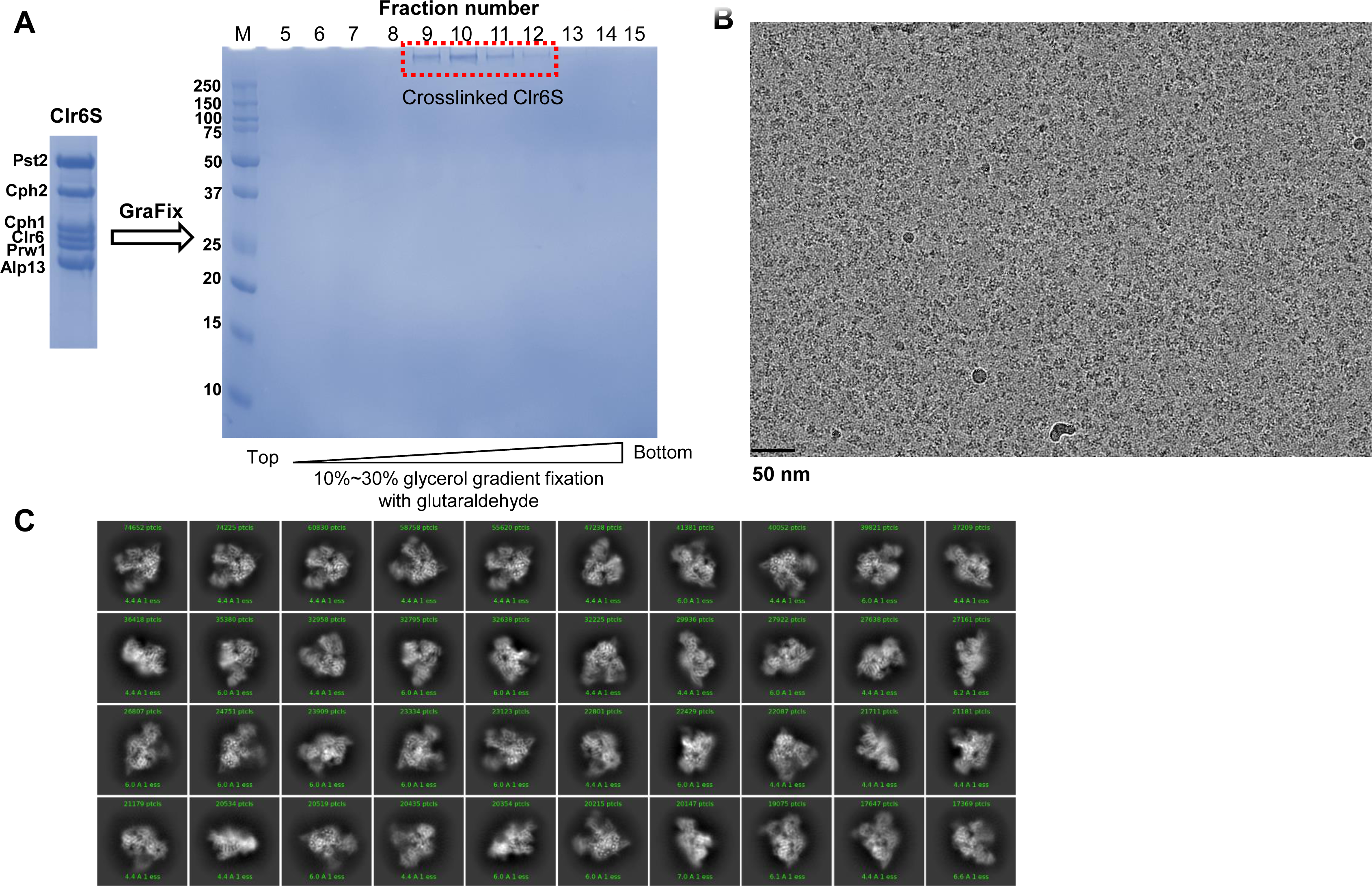
Purification and cryo-EM analysis of Clr6S. (A) SDS-PAGE analysis of the Clr6S fractions after glycerol gradient centrifugation with fixation. The fractions were loaded onto a 12% resolving gel and stained with Coomassie blue. The peak fractions selected for cryo-EM study are indicated with a red dashed square. The fraction numbers are labeled on the top. (B) A representative electron micrograph of Clr6 complex. Scale bar, 500 Å. (C) Representative 2D class averages of Clr6S particles.

**Fig S2.**
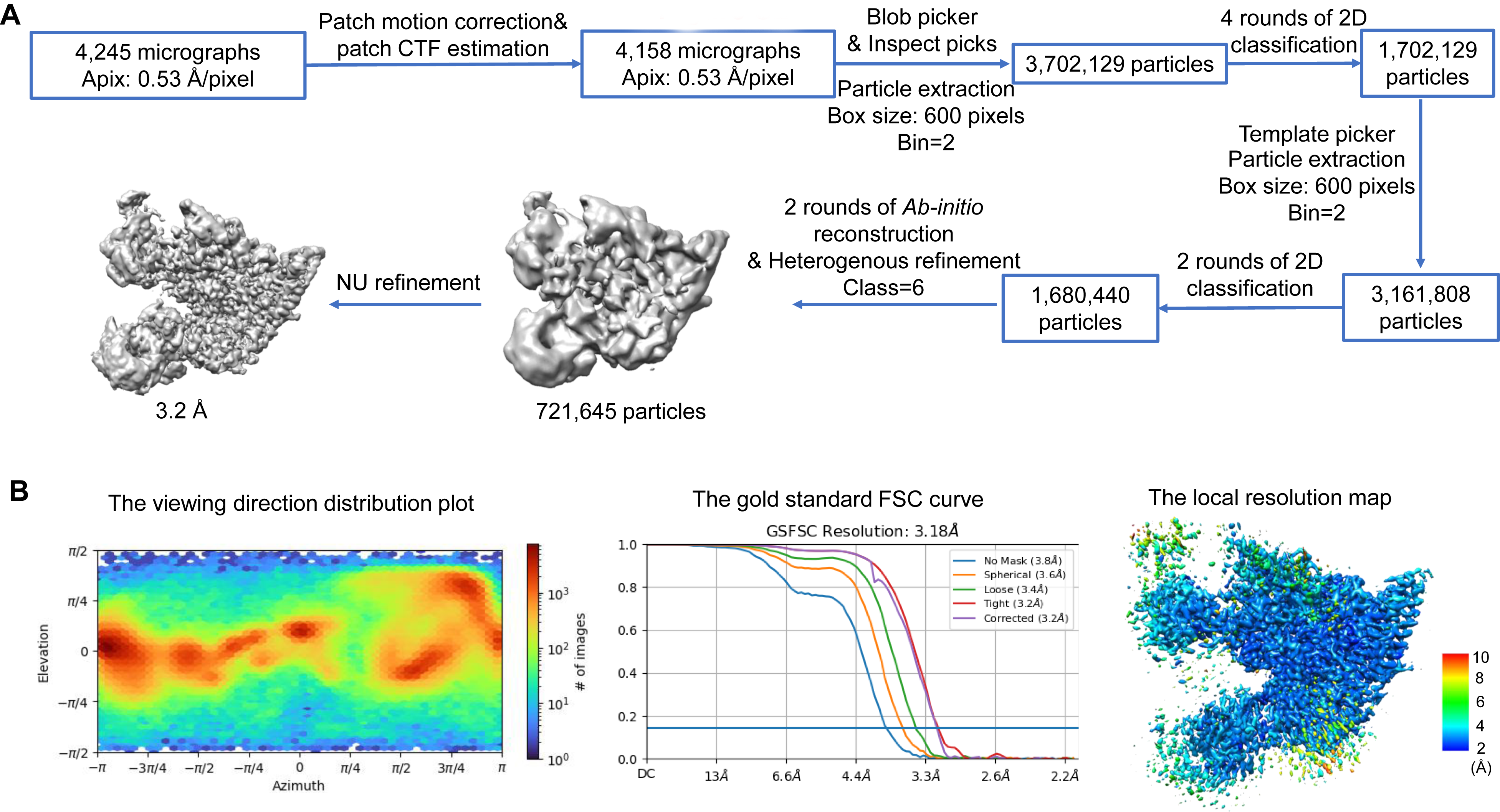
Cryo-EM analysis of Apo-Clr6S. (A) Flow-chart of cryo-EM data processing of the Apo-Clr6S complex. (B) Quality of the cryo-EM maps. From left to right are the viewing direction distribution plot, the 0.143 gold standard FSC curve, and the color-coded local resolution map, respectively.

**Fig S3.**
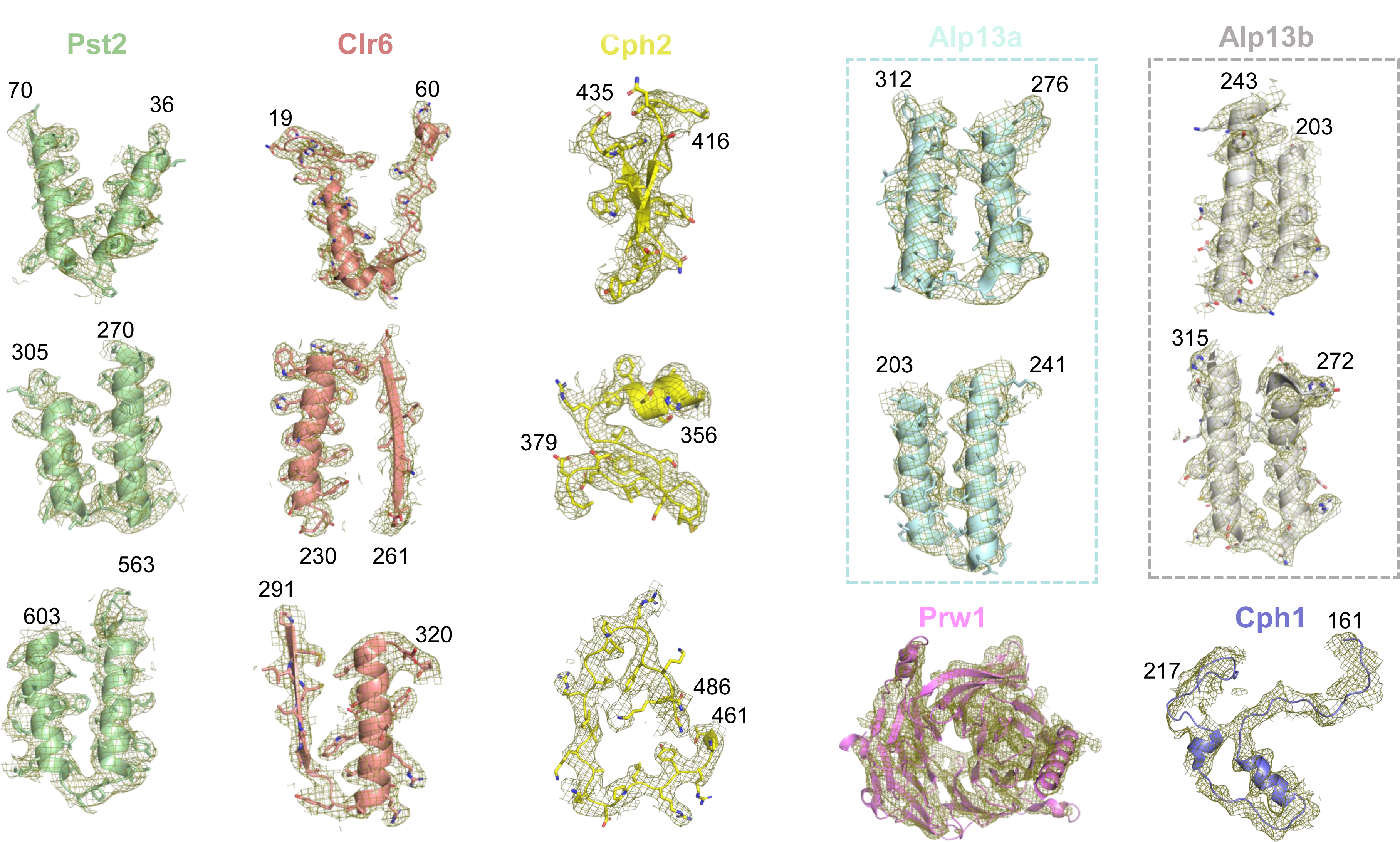
Fitting the models of Pst2, Clr6, Cph1, Cph2, Alp13a, Alp13b and Prw1 to the actual EM density map. Fittings were done in PyMOL. The EM density for representative regions of each subunit is shown in mesh. The amino acids of two ends are labeled.

**Fig S4.**
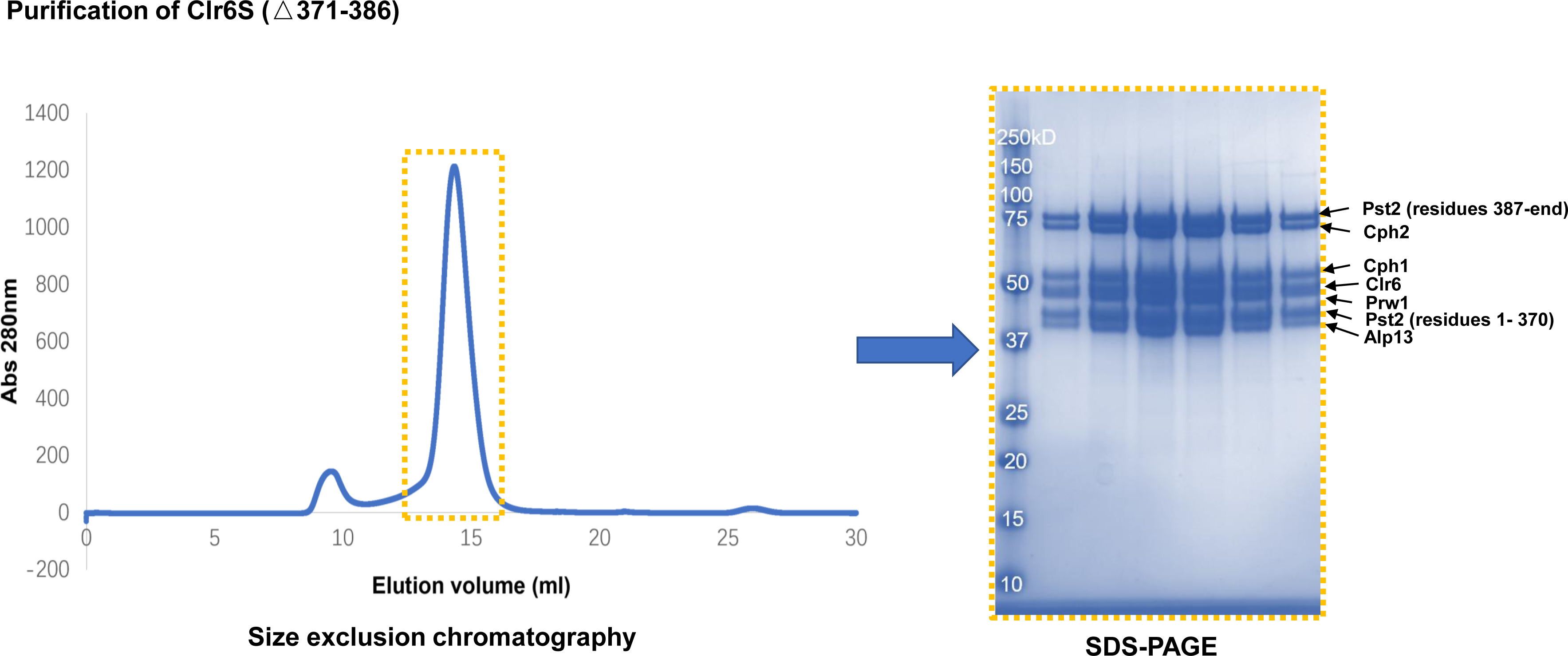
Purification of Clr6S (△371-386). (A) Size exclusion chromatography of Clr6S (△371-386). The peak fractions selected for further SDS-PAGE analysis are boxed with an orange dashed frame. (B) SDS-PAGE analysis of the peak fractions in Fig. S4A. Each band is indicated with a black arrow and labeled on the right.

**Fig S5.**
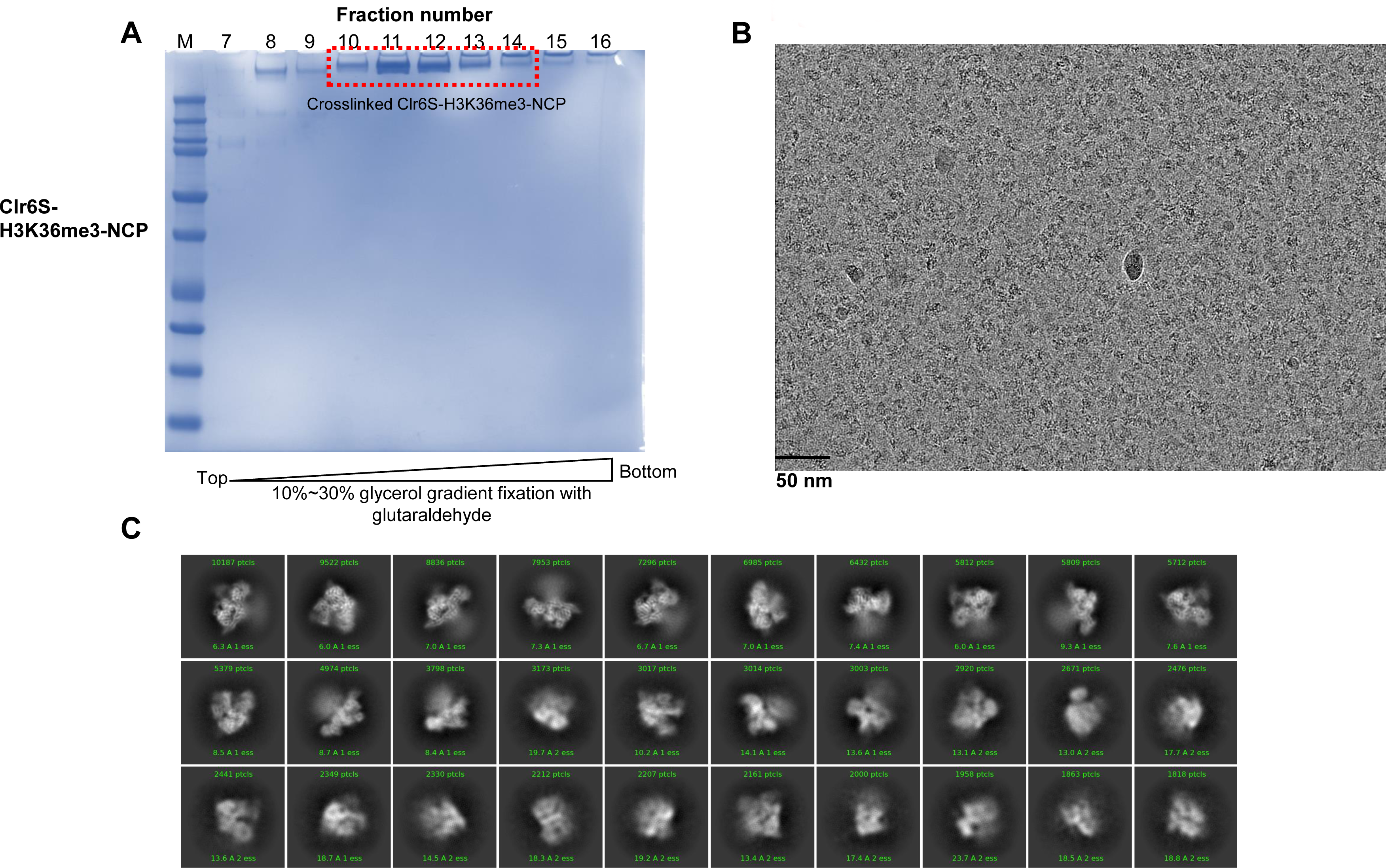
Purification and cryo-EM analysis of Clr6S-H3K36me3-NCP. (A) SDS-PAGE analysis of the Clr6S-H3K36me3-NCP fractions after glycerol gradient centrifugation with fixation. The fractions were loaded onto a 12% resolving gel and stained with Coomassie blue. The peak fractions selected for cryo-EM study are boxed and the fraction numbers are labeled on the top. (B) A representative electron micrograph of Clr6-H3K36me3-NCP complex. Scale bar, 500 Å. (C) Representative 2D class averages of Clr6S-H3K36me3-NCP particles.

**Fig S6.**
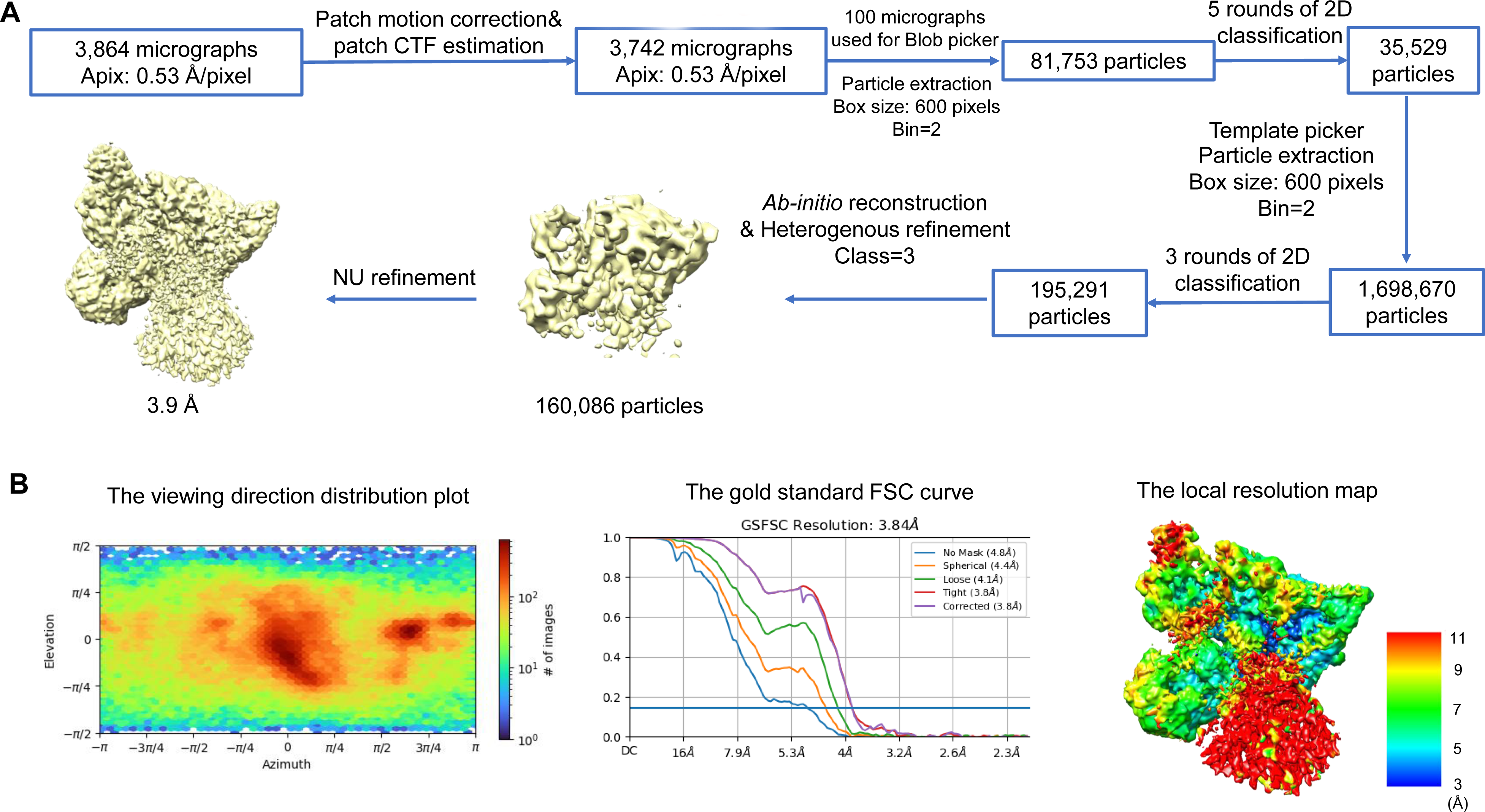
Cryo-EM analysis of Clr6S-H3K36me3-NCP. (A) A flow-chart of cryo-EM data processing of the Clr6S-H3K36me3-NCP. (B) Quality of the cryo-EM maps. From left to right are the viewing direction distribution plot, the 0.143 gold standard FSC curve, and the color-coded local resolution map, respectively.

**Fig S7.**
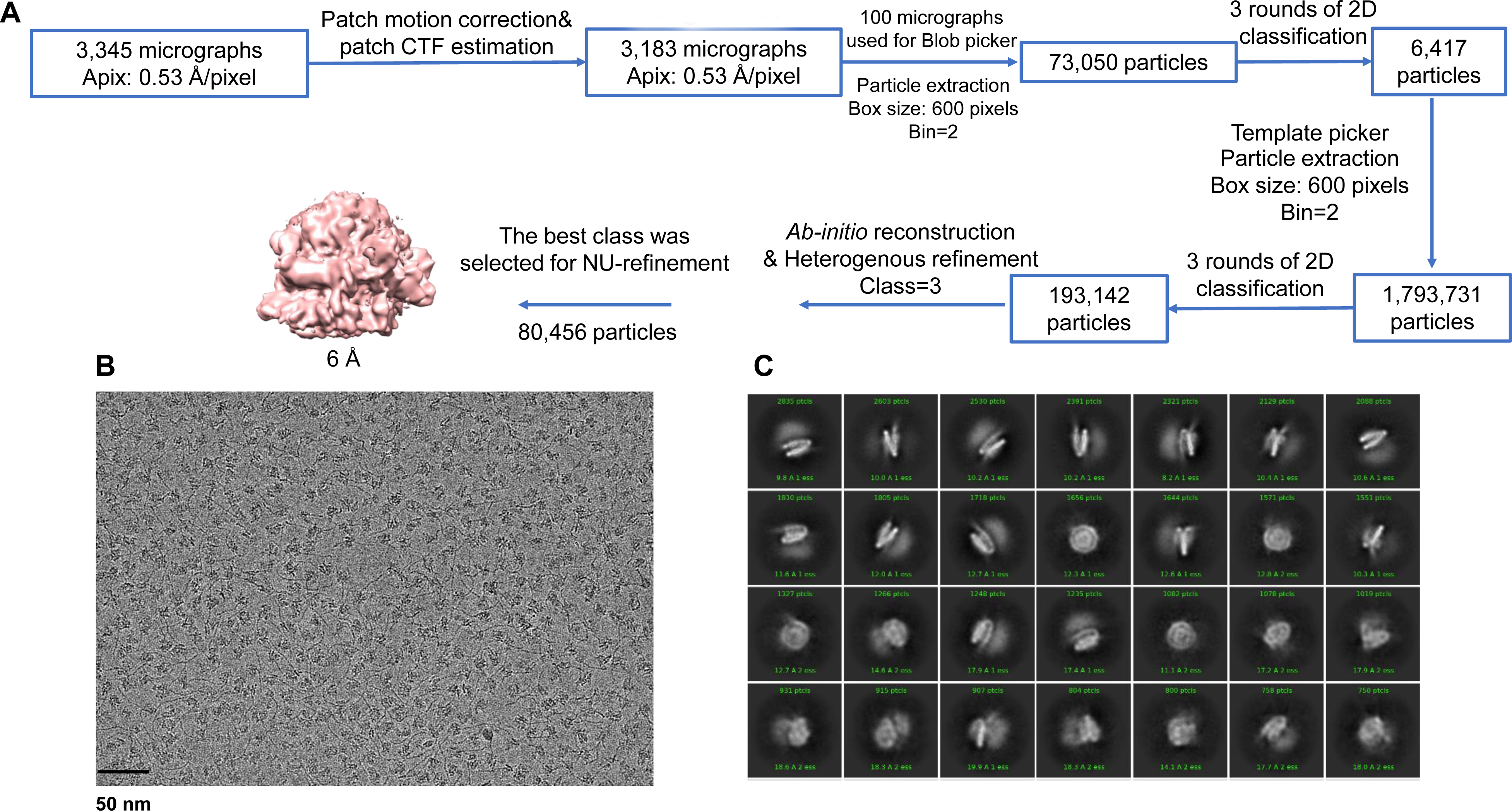
Cryo-EM analysis of the Rpd3S-NCP complex. (A) A flow-chart of cryo-EM data processing of the Rpd3S-H3K36me3-NCP. (B) A representative electron micrograph of Rpd3S-NCP complex. Scale bar, 500 Å. (C) Representative 2D class averages of Rpd3S-NCP particles.

**Fig S8.**
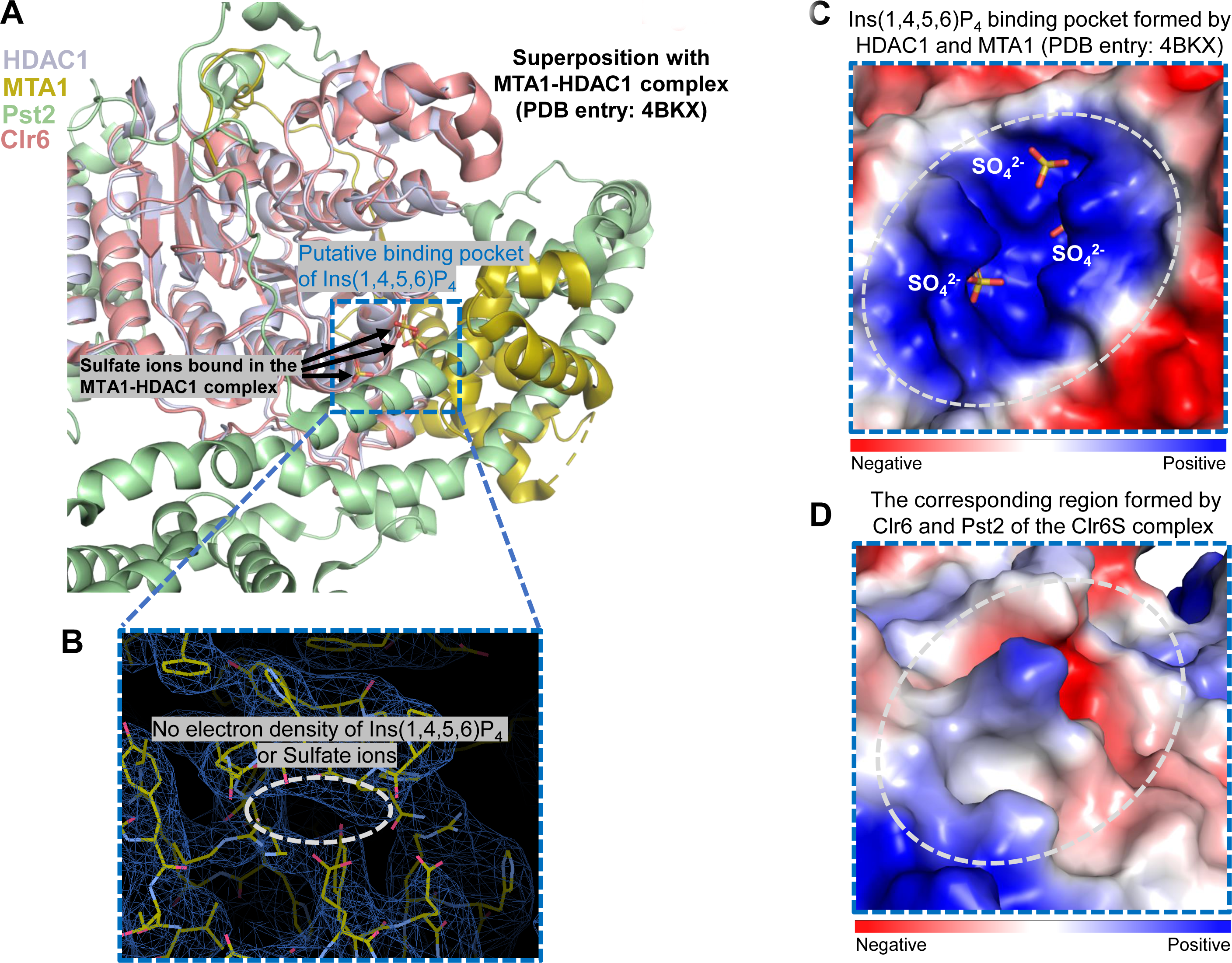
Comparison of the structures of Clr6S and HDAC1-MTA. (A) Structural superposition of Clr6S and HDAC1-MTA from the NuRD complex (PDB entry: 4BTX). The structures of different proteins are color-coded: Clr6 (salmon), Pst2 (light green), HDAC1 (blue white), and MTA1 (sand). The Ins(1,4,5,6)P_4_ binding pocket of HDAC1-MTA1 is highlighted with a dashed blue box. (B) The electron density of the corresponding region in the Clr6S complex. The potential Ins(1,4,5,6)P_4_ binding site is indicated with a dashed white ellipse (no electron density was observed). The figure was generated using COOT. (C) and (D) Comparison of the electrostatic surface of the Ins(1,4,5,6)P_4_ binding pocket in the HDAC1-MTA complex and the corresponding region in the Clr6S complex. The sulfate ions bound in the HDAC1-MTA1 complex are shown in stick representation. Top: the positively charged surface of the Ins(1,4,5,6)P_4_ binding pocket in the HDAC1-MTA1 complex is circled with a dashed white ellipse. Bottom: the electrostatic surface (corresponding to the Ins(1,4,5,6)P_4_ binding pocket in HDAC1-MTA1) on the Clr6S complex is circled with a dashed white ellipse.

