## Supplemental Table 1 for "Class I histone deacetylase complex: structure and functional correlates"

**Table S1. Cryo-EM data collection, refinement, and validation statistics.**

| Complex | Clr6S | Clr6S-H3K36me3-NCP | Rpd3S-H3K36me3-NCP |
|---|---|---|---|
| EMDB | EMD-35416 | EMD-35417 | EMD-XXXXX |
| PDB | 8IFG | | |
| **Data collection and processing** | | | |
| Magnification | 22,500 | 22,500 | 22,500 |
| Voltage (kV) | 300 | 300 | 300 |
| Electron exposure ($e^-/Å^2$) | 60 | 60 | 60 |
| Defocus range (-μm) | 1.3-2.5 | 1.3-2.0 | 1.3-2.0 |
| Pixel size (Å) | 0.53 (super-resolution mode) | 0.53 (super-resolution mode) | 0.53 (super-resolution mode) |
| Symmetry imposed | C1 | C1 | C1 |
| Initial particle images (no.) | 3,161,808 | 1,698,670 | 1,793,731 |
| Final particle images (no.) | 721,645 | 160,086 | 80,456 |
| Map resolution (Å) | 3.2 | 3.9 | 6.0 |
| FSC threshold | 0.143 | 0.143 | 0.143 |
| Map resolution range (Å) | 2.5-3.5 | | |
| **Refinement** | | | |
| Initial model used | AlphaFold2-predicted | | |
| Model resolution (Å) | 3.2 | | |
| FSC threshold | 0.143 | | |
| Model resolution range (Å) | 2.5-3.5 | | |
| Map sharpening B factor ($Å^2$) | 132.6 | | |
| Model composition | | | |
| Non-hydrogen atoms | 19,068 | | |
| Protein residues | 2,338 | | |
| Ligands | 0 | | |
| B factor ($Å^2$) | | | |
| Protein | 167.2 | | |
| Ligand | - | | |
| R.m.s. deviations | | | |
| Bond lengths (Å) | 0.008 | | |
| Bond angles (°) | 0.738 | | |
| Validation | | | |
| MolProbity score | 2.68 | | |
| Clashscore | 13.2 | | |
| Poor rotamers (%) | 0.4 | | |
| Ramachandran plot | | | |
| Favored (%) | 91.4 | | |
| Allowed (%) | 8.4 | | |
| Disallowed (%) | 0.2 | | |
